# MANTIS: Analytics toolkit for spatial metabolomics with matching spatial transcriptomics data

**DOI:** 10.64898/2026.01.20.700581

**Authors:** Yu Hao, Yeojin Kim, Bhavay Aggarwal, Saurabh Sinha

**Author notes:** Corresponding Author Saurabh Sinha.

## Abstract

**Motivation:** Joint Spatial Metabolomics (SM) and Spatial Transcriptomics (ST) profiling is a powerful approach to fine-mapping of metabolic states associated with tissue function. Current computational tools for analysis of “SM+ST” data focus primarily on alignment and integration of the two modalities, with limited support for probing biological relationships between the two molecular layers.

**Results:** We present MANTIS, a statistical framework for analyzing co-registered SM+ST profiles at single cell or spot resolution, along with spatial domain or cell type information, to discover metabolite spatial patterns and gene-metabolite relationships. It employs an autocorrelation-preserving permutation strategy to assess statistical significance, yielding calibrated inference under spatial dependence. It disentangles different sources of spatial patterns and correlations, viz., those arising from regional preferences, cell type associations, or other unknown factors. It introduces the use of spatial cross-correlation and spatial partial correlation statistics for quantifying gene-metabolite associations. Across data sets spanning different spatial technologies, tissues and species, MANTIS provides more specific and interpretable discoveries than existing methods through rigorous statistical testing and explicitly modeling confounding structure. To our knowledge, MANTIS is the first toolkit to unify spatial metabolomics, spatial transcriptomics, cell type information and spatial domains within a single framework that emphasizes spatial statistics, hypothesis testing and confounder correction.

**Availability and Implementation:** Freely available on the web at https://github.com/yuhaotuo/MANTIS.

## INTRODUCTION

The rapidly maturing technology of spatial metabolomics [1], where abundances of many metabolites are measured in a multiplexed manner with spatial resolution, can be transformational for fine-mapping of cellular processes associated with tissue function. For instance, spatial metabolomics (SM) maps of the brain reveal lipid- and metabolite-level variations associated with function in normal [2] as well as disease conditions [3,4]. Recent breakthroughs have enabled SM profiling and spatial transcriptomics (ST) profiling of the same tissue, providing an even richer multi-omics view into the tissue’s molecular state [5]. The matched ST profile can help delineate cell types and spatial domains [6] in the tissue, giving us clarity on the regional selectivity [7] and cell type-specificity of metabolic events and their changes. For instance, Hu et al. [1] used matched SM and spatial proteomics profiling to characterize local metabolite competition of neighboring cells in human lung cancer. Similarly, Vicari et al. [5] analyzed SM+ST profiles of mouse striatum to characterize genes and neuronal subtypes associated with dopamine. The ST profile can also elucidate regulatory mechanisms related to metabolic alterations [8].

The transformative potential of joint metabolomics and transcriptomics profiling cannot be unlocked without the key to integrative multi-omics *analysis*. Spatial metabolomics analysis lacks comprehensive statistical tools for unbiased discovery of metabolic patterns involving cell types and spatial domains. Existing tools for SM data offer a variety of core functionalities such as annotating m/z peaks, dimensionality reduction, clustering and visualization, spatial structure discovery, pairwise metabolite network reconstruction, etc. [2,9,10]. However, these tools operate at the *pixel-level* rather than cell-level, and typically do not incorporate cell type or spatial domain information. Recently introduced tools tackle SM+ST data, addressing cross-modality alignment and cell segmentation [2,11], a necessary step for cell-level analysis. SpaMTP [12] is a state-of-the-art tool for downstream analysis of such cell-level multi-omics data, including gene-metabolite correlation analysis, which can provide important clues into mechanisms associated with metabolic patterns. SpatialMETA [13] uses neural networks for cross-modality (SM, ST) integration and provides several useful downstream analyses of the integrated data, including spatial clustering, markers of clusters, gene-metabolite correlation, etc. However, key analytic goals remain unmet in existing tools for SM+ST data analysis. For instance, current tools [12,13] for discovering gene-metabolite relationships use non-spatial correlation metrics such as the Pearson correlation coefficient; ideally such relationships should be quantified in a spatially aware manner, in order to account for prevalent and diverse spatial covariance structures that may confound the findings. Gene-metabolite correlations may also be confounded by associations with cell types and regions, and careful accounting of these confounders is necessary. Furthermore, correlation scores need to be accompanied by robust statistical testing in order to control false discoveries, an aspect that has not been rigorously addressed in prior work. Finally, while existing tools do provide some support for detecting region- or cell type-associated metabolites, such functions are unable to disentangle spatial variation driven by tissue organization, cell type distributions and other factors, or to assess statistical significance of these effects in a unified framework.

Here, we present a computational toolkit called **MANTIS (M**etabolomics **An**d **T**ranscriptomics **I**n **S**pace) for analysis of paired spatial metabolomics and transcriptomics (SM+ST) data. It provides functions for two broad categories of tasks: (1) discovery of metabolic spatial patterns and (2) identification of gene-metabolite relationships. It exploits cell type annotations of cells and spatial domain delineations of the tissue, if available, to extract biologically interpretable patterns and relationships. (See Table 1.) It employs well-established spatial and non-spatial statistical approaches and a novel strategy to assess statistical significance of spatial patterns. It also disentangles different sources of spatial patterns and correlations, viz., those arising from regional preferences or cell type associations of a metabolite. We present the functions of the toolkit with examples of their application to three data sets where SM profiles are supported by ST profiles of the same or adjacent tissue section. The SM datasets used in this study were generated using different technologies, illustrating the ability of MANTIS to operate across heterogeneous measurement platforms. To our knowledge, MANTIS is the first software toolkit to combine spatial, single-cell metabolomics, transcriptomics, and cell type information with rigorous statistical procedures, revealing metabolite patterns and their gene-level correlates and mechanisms.

|  | Feature | MANTIS | SpatialMETA | SpaMTP |
| --- | --- | --- | --- | --- |
| Metabolite Analysis | Cell Type-associated metabolite discovery | ✓ | ✓ |  |
|  | Region-associated metabolite discovery | ✓ | ✓ | ✓ |
|  | Spatially variable metabolite discovery adjusted for cell type or region | ✓ |  |  |
| Metabolite-gene association | Statistical method | Spatial correlation | PCC |  |
|  | Adjusted for cell type- or region-associations | ✓ |  |  |
| Spatially aware null model for significance testing (autocorrelation-controlled) |  | ✓ |  |  |
| Metabolite annotation |  |  | ✓ | ✓ |
| Alignment |  |  | ✓ | ✓ |
| Clustering |  |  | ✓ | ✓ |

## RESULTS

### Overview of MANTIS toolkit

MANTIS provides a suite of functions for analyzing spatial metabolomics (SM) data on a tissue sample along with co-registered spatial transcriptomics (ST) data for the same or adjacent tissue section. To account for variations in spatial resolution across technologies, we assume that the data set comprises a set of “objects” with a metabolomics profile, a transcriptomics profile and spatial coordinates (**Figure 1**). Objects may be individual cells, or “spots” in certain ST technologies such as 10x Visium. We assume that mismatches in resolution have been addressed and objects are aligned [5,12], for instance through available tools such as SpatialMETA [13] and SpaMTP [12]. We also assume that the ST data have been analyzed via third-party tools to annotate cell types [14] or to deconvolve spots into cell type proportions [15] and to delineate spatial domains of the tissue [14]. Several tools are available for both tasks [14], and studies generating ST data typically provide cell type annotations and often spatial domains (e.g., brain regions [16], tumor clusters and microenvironments [1], etc.) Thus, the input to MANTIS can be described as in **Figure 1A**, as a matrix with rows representing “objects” (cells, spots, etc.), and columns representing information about those objects, e.g., spatial coordinates, metabolite abundances and gene expression, and optionally region and/or cell type annotations. MANTIS includes a set of functions that analyze such a data matrix to report lists of metabolic spatial patterns and gene-metabolite associations (Figure 1), with rigorous significance assessment and handling of confounding factors.

**Figure 1.**
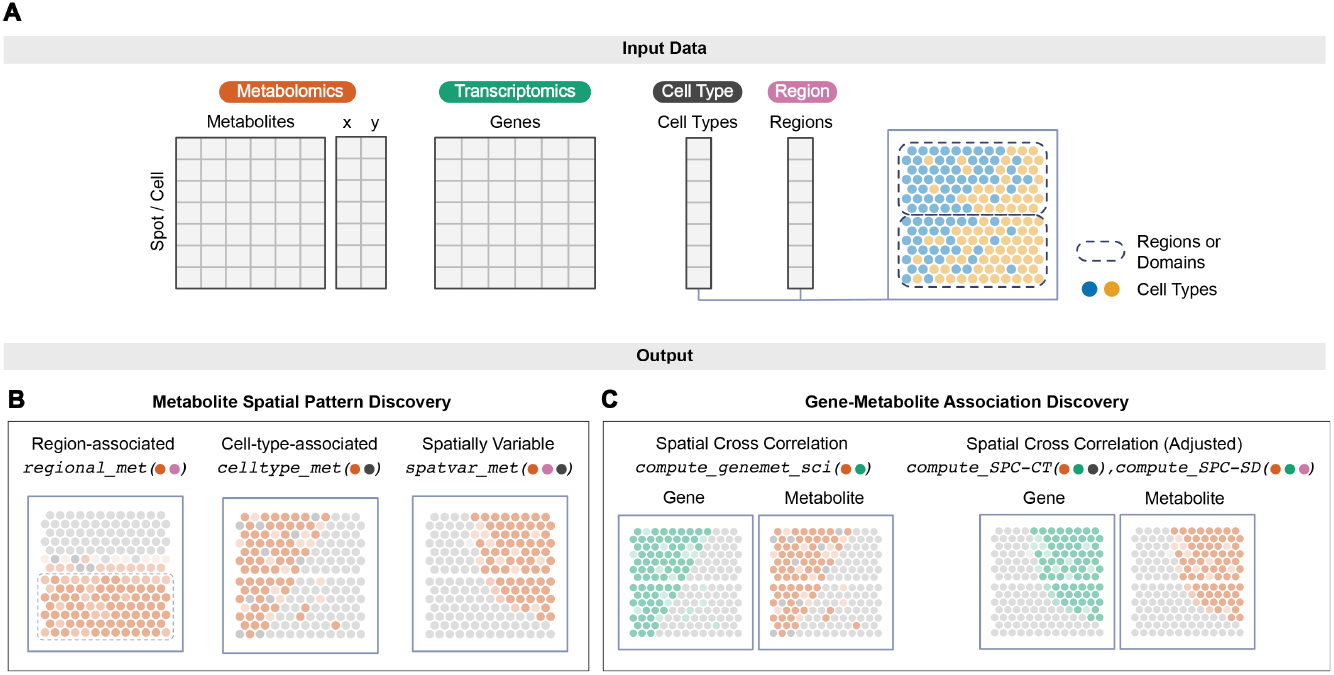
Overview of MANTIS. **(A)** MANTIS analyzes spatial multi-omics data profiling metabolite intensities and gene expression levels for each cell or spot in a tissue section. The input is assumed to be metabolomics profiles and transcriptomics profiles of cells/spots along with their spatial locations. Optionally, the cells/spots may be annotated with their spatial domains (outlined by dashed lines) and/or cell types (colored circles), to be used to decompose sources of spatial variation. **(B)** One set of routines in MANTIS detect metabolites with interesting spatial patterns, including associations with spatial domains (regions) or cell types, as well as non-random spatial patterns not easily attributed to regions or cell types. **(C)** Another set of routines report gene-metabolite pairs with correlated spatial patterns (gene shown in left panel, metabolite in right panel).

We seek to identify metabolites with “interesting” (statistically significant) spatial distributions. Metabolomic mapping studies have pointed to many such metabolic spatial patterns in mouse brain [17], human and primate brains [18], human tonsil and tumor samples [1], etc. Of interest is a metabolite that is (1) abundant in one or a few anatomical/functional regions of the tissue, (2) is expressed in cellular neighborhoods (“niches”) marked by specific cell types, or (3) exhibits a statistically unusual spatial distribution even if it does not segregate by regions/cell types [19] (**Figure 1B**). Such patterns may help us link metabolites to regional functions of the tissue [17], reveal metabolic requirements or signaling roles of specific cell types [17,20], or point to previously unrecognized metabolic microenvironments or functional domains within the tissue [2,20-22]. MANTIS provides functions to identify these three classes of metabolite spatial patterns. In addition, MANTIS provides routines to quantify gene-metabolite associations via spatial statistics (Figure 1C), while also accounting for potential confounders such as regions and cell types. We illustrate these functions via application to the mouse brain striatum SM+ST data set of Vicari et al. [5], referred to below as the VICARI data set.

### Discovery of region-associated metabolites

A simple and easily interpreted pattern is that of a metabolite preferentially localized to one or more spatial domains (“regions”). Regional preference of metabolites often reflects specialized functions of brain areas, e.g., certain neurotransmitters enriched in regions active in signaling, or energy metabolites concentrated in areas with high metabolic demand [21]. MANTIS provides the ‘**regional_met**’ function to identify such patterns. This function first represents a metabolite’s abundances across regions as a probability vector: the “regional abundance profile” for that metabolite. A background profile is obtained similarly by averaging abundances of *all* metabolites in each region. It then quantifies the divergence between the metabolite’s profile and background profile using Kullback-Leibler (KL) divergence (**Methods**). This is repeated for every metabolite, and significance threshold is set based on an empirical null distribution obtained by a custom permutation method that creates randomized versions of a metabolite’s spatial distribution while preserving autocorrelation, see **Methods**.

Using the regional_met function on the VICARI data set, which includes 9 delineated spatial domains (regions in the mouse striatum) (**Figure 2A**) we obtained a list of 66 significant metabolic spatial patterns (FDR < 0.1, **Figure 2D, Supplementary Table S1**) out of the 500 highly variable metabolites examined. **Figures 2B and 2C** show two such metabolites – m/z 725.551, which is enriched in the ventrolateral striatum (VL), and m/z 822.614, which is relatively abundant in the corpus callosum (CC) and anterior commissure (ACO). The complete set of region-associated metabolites spans a small set of patterns (**Figure 2E, Supplementary Figure S1**), several of which are similar to cell types’ distributions (**Supplementary Figure S2**), suggesting that metabolite-cell type associations may underlie their regional preferences (or vice versa).

**Figure 2.**
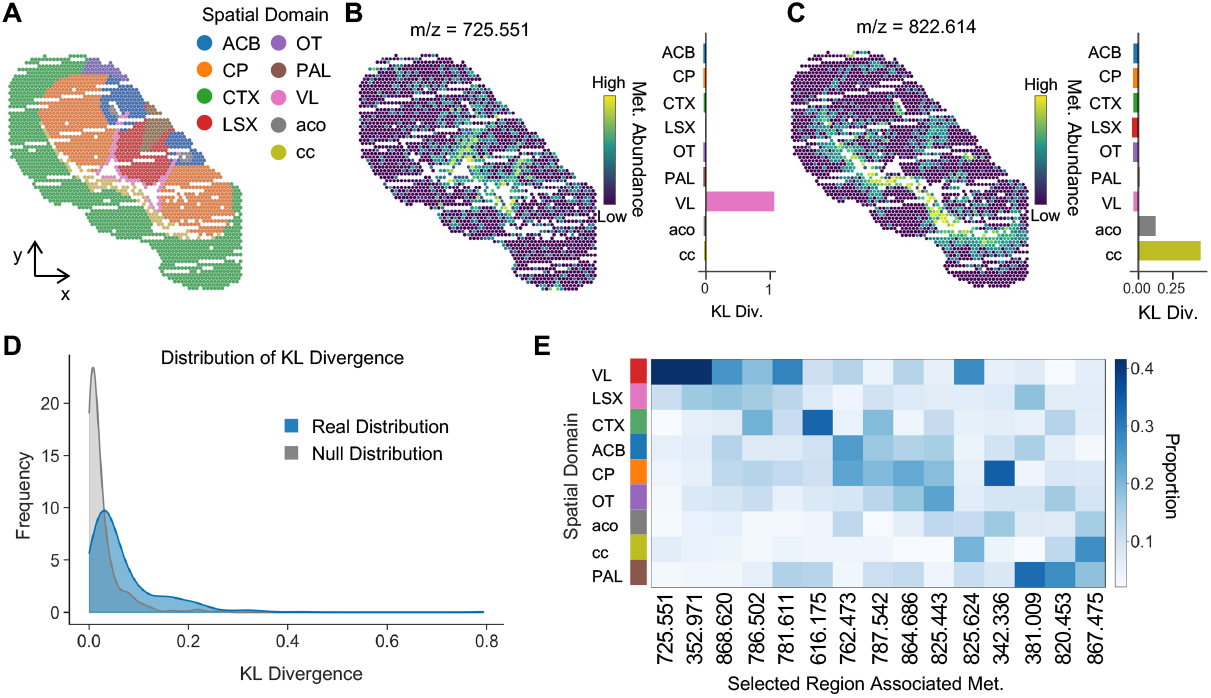
Region-associated metabolites in mouse striatum SM+ST data (VICARI). **(A)** Spatial domains of the striatum profiled in the data set. ACB: Nucleus accumbens, CP: Caudate putamen, CTX: Cortex, LSX: Lateral septal complex, OT: Olfactory tubercle, PAL: Pallidum, VL: Ventral striatum, aco: anterior commissure, cc: corpus callosum. **(B, C)** Examples of region-associated metabolites discovered by MANTIS. Left subpanels show metabolite abundance maps; right subpanels show the metabolite’s regional abundance profile and its enrichment (log fold change relative to average of all metabolites) in specific domains. m/z 725.551 is enriched in VL (B), while m/z 822.614 shows preferential abundance in cc/aco (C). **(D)** Distribution of KL-divergence scores of metabolites, representing their statistical association with spatial domains. The observed distribution over 500 metabolites in the VICARI data (blue) is contrasted with an empirical null distribution (gray) constructed with a randomization procedure that preserves spatial autocorrelations. **(E)** Heatmap showing regional abundance profiles of select metabolites. Color intensities represent abundance of a metabolite (column) in each region (row). Shown metabolites were selected to illustrate major patterns of region-associations seen across the complete set of significant region-associated metabolites.

*Comparison to existing tools:* Available tools such as SpatialMETA and SpaMTP provide a similar functionality of identifying “marker” metabolites, by testing if a metabolite’s abundance is enriched in a region compared to other regions, using a between-group difference test [13]. Our approach is different in that (a) it tests for non-randomness of the overall regional distribution of a metabolite rather than enrichment in each region separately, thus avoiding a multiple testing burden, and (b) performs a statistical test that controls for the spatial autocorrelation of the metabolite, which is not addressed in current tools. We show in **Supplementary Table S2** that the “one region vs. others” test reports a very large number of significant associations. For instance, a Wilcoxon test finds 2,886 metabolite-region associations at FDR < 0.05 (64% of 4,500 pairs tested). In contrast, MANTIS performs only 500 tests (one per metabolite) and found 66 significant region-associated metabolites at FDR < 0.1. Though there is no “ground truth” to objectively establish their relative error rates, we interpret these findings as suggesting that MANTIS reveals a more conservative set of regional associations that is likely to have a lower false positive rate.

### Discovery of cell type-associated metabolites

There have been several reports of specific metabolites being enriched in certain cell types and in tissue regions with specific cell type compositions [1,17,20]. These associations may be specific to a region, e.g., elevated glycogen in astrocytes in the hippocampus and cerebellar cortex [23], or transcend regional boundaries, e.g., lactate predominantly released by astrocytes to fuel neuronal activity [24]. Findings of metabolite-cell type associations can point to instances of metabolic reprogramming in certain disease contexts.

MANTIS provides the ‘**celltype_met**’ function to discover cell type-associated metabolites. It tests for spatial covariance between a cell type’s presence and a metabolite’s abundance, using the Spatial Cross-Correlation Index (SCI) [25] (**Methods**), which is closely related to bivariate Moran’s I [26]. For spot-level data, cell type presence is assumed to be obtained from deconvolution tools while for cell-level data a Boolean indicator of cell type annotation is used. A permutation-based empirical null distribution that controls for metabolite spatial autocorrelations is used to set a significance threshold on the SCI score (**Methods**).

Using this function on the VICARI data set, which contains 48 different cell types (**Figure 3A**), we obtained 1,518 significant cell type-metabolite associations (FDR<0.05) (**Supplementary Table S3**), with the most associations being for oligodendrocytes (OLIG), Schwann Cells (SCHW) and Satellite glia (SATG) (**Figure 3B**). **Figure 3C** depicts the strong spatial correlation between cell type Cerebellum neurons and metabolite 172.001 m/z (SCI score 0.474). Since certain cell types are statistically segregated across spatial domains, we expected and confirmed that many of the discovered cell type-metabolite associations are also region-associated metabolites (**Figure 3D**).

**Figure 3.**
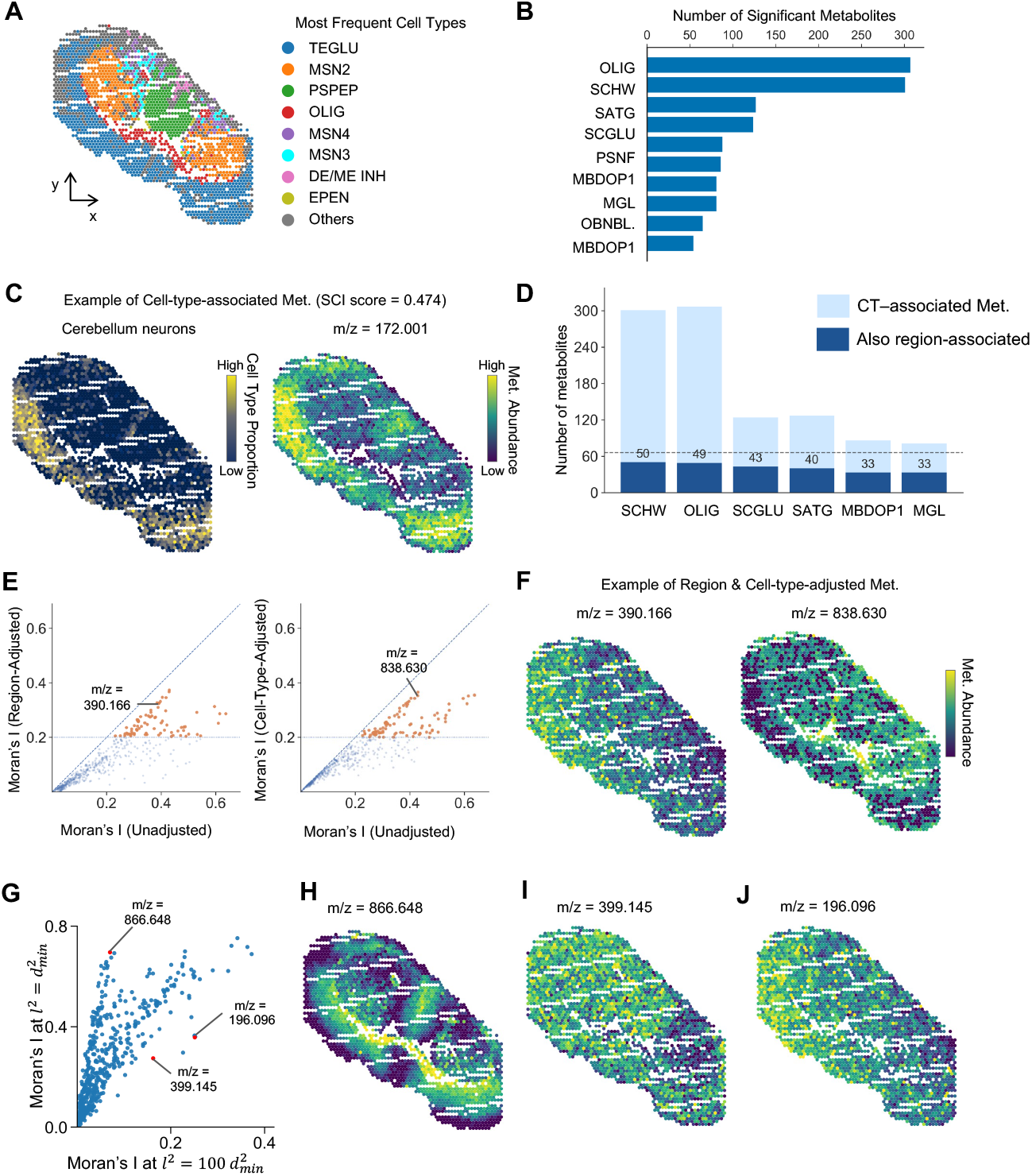
Cell-type-associated metabolites and spatially variable metabolites in VICARI data. **(A)** Spatial distribution of eight most frequent cell types in the mouse striatum section. TEGLU (Telencephalic Glutamatergic Neurons), MSN2 (Medium Spiny Neurons, Subtype 2), PSPEP (Pallidal/Striatal Peptidergic Neurons), OLIG (Oligodendrocytes), MSN4 (Medium Spiny Neurons, Subtype 4), MSN3 (Medium Spiny Neurons, Subtype 3), DE/ME INH (Diencephalic/Mesencephalic Inhibitory Neurons), EPEN (Ependymal Cells), and Others (other cell-types). **(B)** Counts of significant cell-type-metabolite associations reported by MANTIS, for each cell type; OLIG, SCHW, and SATG show the largest numbers of associations. **(C)** Example of significant cell-type-metabolite association: Cerebellum neurons proportion (left) and metabolite m/z 172.001 (right) show a high SCI score of 0.474. **(D)** Cell-type-associated metabolites often include region-associated metabolites, as cell types themselves exhibit regional preferences. For each cell type, the bar shows the number of metabolites associated with that cell type and the number of such metabolites that are also region-associated. The dashed line represents the total number of region-associated metabolites (n = 66). **(E)** Effect of confounder adjustment for spatial autocorrelation statistics (Moran’s I) used to detect spatially variable metabolites (SVMs). Scatter plots compare Moran’s I of each metabolite’s spatial distribution (unadjusted, x-axis) vs adjusted values after regressing out region (left) or cell-type (right). **(F)** Examples of metabolites retaining spatial autocorrelation signal (high Moran’s I) after adjustment for region and cell type; the unadjusted vs adjusted scores for these two examples are shown in panel E. **(G)** Effect of the “length scale” parameter used in Moran’s I. Scatter plot compares Moran’s I at two extreme spatial scales (*l* = *d*_*min*_ and *l* = 100*d*_*min*_), and two examples, m/z = 866.648, m/z = 399.145 and m/z = 196.096, are highlighted to illustrate opposite trends. **(H)** Feature map of m/z = 866.637, which exhibits strong fine-grained boundaries and high Moran’s I at small *l* but loses spatial autocorrelation at larger *l*. **(I-J)** Feature maps of m/z = 399.145 and m/z = 196.096, which display a broad gradient that becomes more obvious at large *l*.

#### Comparison to existing tools

To our knowledge, existing tools do not explicitly test for cell type associations of metabolites. A possible baseline approach is to use Pearson correlation or Wilcoxon test depending on whether cell type information is quantitative (proportions) or binary. Such an approach differs from MANTIS in two ways: the score is calculated in a spatially agnostic manner, and the significance test does not account for spatial autocorrelations. We evaluated this baseline method (**Supplementary Table S4**) and found 11,869 significant cell type-metabolite associations at FDR < 0.05, ∼49.4% of 24,000 tested pairs. In the absence of a reliable “ground truth” we are unable to perform an objective comparison of error rates to that of MANTIS, but the far larger number of associations reported (at the same FDR threshold) suggests a weak null hypothesis used in the non-spatial baseline. **Supplementary Figure S3** shows an example association that is designated significant by the baseline method but not by MANTIS.

### Discovery of spatially variable metabolites

A metabolite’s spatial pattern may not necessarily segregate by spatial domains or cell type niches but still be relevant to tissue function. We drew inspiration from the literature on “spatially varying genes” (SVGs) [27,28] to search for such patterns in SM data. These tools test if spatially proximal cells tend to exhibit similar levels of the molecule of interest. The ‘**spatvar_met**’ function in MANTIS provides a similar functionality – detection of spatially variable metabolites (SVMs) – using the Moran’s I score [29] . We recognized that a region-associated metabolite is likely to be also detected as a SVM because being enriched in a region implies clustering of cells/spots where the metabolite is found. Similarly, cell type-associated metabolites are also likely to be detected as a SVM. Thus, the spatvar_met function corrects for the confounding effects of spatial domains (or cell type) by first regressing metabolite abundance against spatial domain (or cell type) information and applying the Moran’s I score to the residuals (**Figure 3E, Methods**). A metabolite that retains a high Moran’s I after both adjustments is designated as a spatially variable metabolite.

Using this function on the VICARI data, we discovered 73 spatially variable metabolites (out of 500), based on a Moran’s I threshold of 0.2 (**Supplementary Table S5**). **Figure 3F** shows two distinct examples of such patterns: 390.166 m/z displays a broad gradient emanating from the dorsolateral injection site (dopamine delivered from the upper-left), consistent with diffusion from that source, whereas 838.630 m/z exhibits discrete focal hotspots that do not align well with major domains or local cell-type composition.

The spatvar_met routine includes a user-tunable parameter called the length scale ( *l* ) that influences the type of pattern reported. Smaller *l* increases sensitivity to local structure, while larger *l* accommodates larger-scale patterns. **Supplementary Table S6** shows a smooth, monotonic decrease in the number of SVMs identified with increasing *l*, with **Figures 3G-3I** showing how extreme values of the length scale can reveal complementary spatial patterns. The analyses above (Figures 3E,F) were performed with an intermediate value of 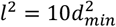 (*d*_*min*_is distance between spots).

#### Comparison to existing tools

The Moran’s I functionality is also provided as part of the SpatialMETA and SpaMTP tools. However, these implementations do not account for the confounding effects of spatial domains and cell types. To underscore the consequences of this difference, we show (**Supplementary Table S5)** that of the 181 (36% of all) metabolites with Moran’s I ≥ 0.2, only 73 retain that level of autocorrelation after accounting for confounders. Thus, the spatvar_met function avoids redundancy and mutual confounding among different types of metabolic spatial patterns. We also tested the SVG detection tool SpatialDE [27] on this data set, identifying all 500 metabolites as significant (FDR < 0.02), most of which can be explained by spatial domain and/or cell type associations, which are not corrected for by that tool (**Supplementary Table S7**).

### Discovering gene-metabolite associations

The routines introduced above primarily analyze SM data and reveal metabolite-associated patterns, using ST data in a supporting role. We now outline the more direct form of cross-omics functionality provided by MANTIS –identifying gene-metabolite pairs with spatially correlated expression. The “**compute_genemet_sci**” function calculates SCI score for every gene-metabolite pair. This score is high if the gene and metabolite exhibit co-expression in the same spots and/or in proximal spots (**Methods**).

We used the compute_genemet_sci function on the VICARI data set, focusing on 1000 highly variable genes and 500 highly variable metabolites (**Supplementary Table S8, Figure 4A**), noting extreme values in excess of 0.6. Visualizations of two sample pairs clearly depicts the high gene-metabolite covariation (**Figure 4B, Supplementary Figure S4**). SCI scores are assessed for statistical significance using an empirical null distribution derived from a randomized data set that preserves the metabolite’s spatial autocorrelation (**Supplementary Figure S5**), and we noted that 12.3% of all gene-metabolite pairs tested were significant at an FDR < 0.05 (Supplementary Table S8).

**Figure 4.**
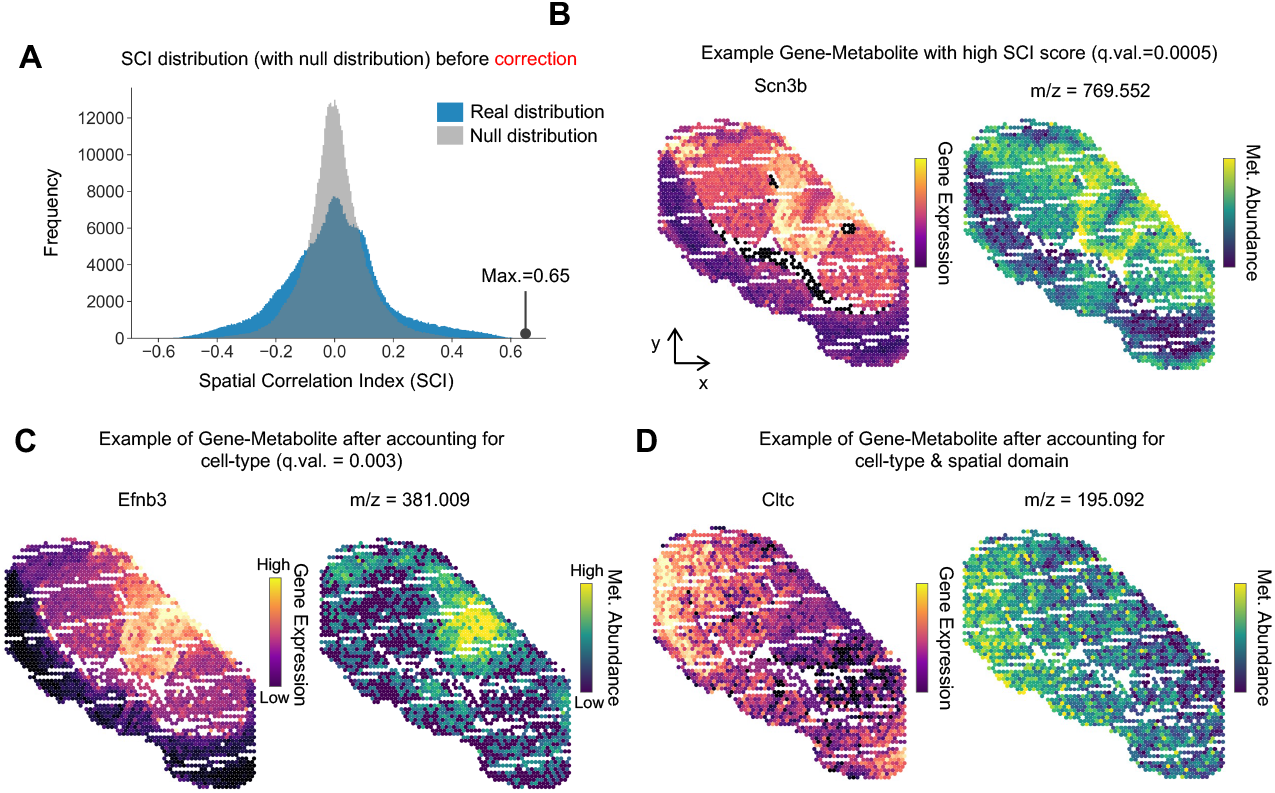
Gene-metabolite spatial covariation. **(A)** Spatial Correlation Index (SCI) scores for all 500,000 gene-metabolite pairs in the VICARI data set (blue), compared to empirical null distribution (gray) derived by a randomization procedure that preserves autocorrelations. **(B)** Example of significant gene-metabolite spatial associations (SCI q-value 0.002). Shown are gene expression of Scn3b (left) and abundance of metabolite 769.552 m/z (right) in the striatum tissue section. **(C)** Example of gene-metabolite pair with significant spatial association (SCI q-value 0.003) even after accounting for their cell type-associations. (SCI is calculated after regressing out the effect of cell type from both variables.) **(D)** Example of a gene–metabolite pair *Cltc* and 195.092 m/z that remains significantly spatially associated even after adjusting for cell-type composition and spatial domain.

A gene and metabolite that are expressed specifically in the same cell type will tend to exhibit high SCI. Furthermore, if there are many genes and metabolites specific to that cell type, then all pairs thereof will be significantly associated. It is more parsimonious to separate out such cases and report them as sets of genes and metabolites associated with that cell type, rather than as directly associated gene-metabolite pairs. Thus, MANTIS provides the “**compute_SPC-CT**” function to report gene-metabolite spatial correlations after removing the confounding effect of cell type information. It uses a statistical score that we call “Spatial Partial Correlation” (SPC), where the gene and metabolite levels are first fit separately to cell type and the residuals of the fit are subjected to SCI calculation (**Methods**).

Using the “compute_SPC-CT” function, we obtained a revised compendium of gene-metabolite associations (**Supplementary Table S9, Supplementary Figure S6**), with 6.8% of the pairs significant at adjusted p-value < 0.05. This confirms that the majority of the significant pairs identified initially (Figure 4A) were likely a result of cell type-specific expression. Two of the strongest gene-metabolite spatial associations observed after accounting for cell type information are shown in **Figure 4C** and **Supplementary Figure S7**. In Figure 4C, the *Raph1*–848.540 m/z pair shows a closely matched spatial pattern across the tissue; this agreement remains after residualizing cell type effects (see **Supplementary Figure S8**). A secondary product of the compute_SPC-CT function is a listing of gene-metabolite pairs whose strong statistical association is primarily driven by their link to cell types (**Supplementary Table S10**). An example of such a pair is shown in **Supplementary Figure S9**.

Another potential confounder for gene-metabolite associations is spatial domains (SD) in the tissue -- a gene and metabolite localized in the same SD are likely to exhibit a high SCI. MANTIS provides the function ‘**compute_SPC-SD**’ to report gene-metabolite associations not explainable by co-localization in a SD **(Methods**) – it first removes SD-related signals from the gene’s and metabolite’s expression distribution and then calculates SCI on the residuals. Applying this function yielded 9.4% of the tested gene-metabolite pairs as significantly associated at adjusted p-value < 0.05 (**Supplementary Table S11**). Intersecting these pairs with those from **Supplementary Table S9**, we obtained a list of 11,890 (2.4% of the 500,000 tested) gene-metabolite pairs whose associations are not attributable to co-expression in certain cell types or spatial domains (**Supplementary Table S12, Figure 4D**).

#### Comparison with state-of-the-art tools

The only available functionality for detecting gene-metabolite relationships in SM+ST data is the correlation routine in SpaMTP and SpatialMETA based on Pearson correlation coefficient (PCC). To assess the difference between this and the SCI-based approach of MANTIS, we calculated PCC between every gene-metabolite pair, observing ∼385K pairs (77% of the 500K tested) to be significant at FDR < 0.05 (**Supplementary Table S13**). This is a far greater number than the 61,552 (12.3% of 500K tested) pairs reported by the compute_genemet_sci routine (above). The wide gap is attributable to the use of a spatial cross-correlation score (SCI) in place of the standard correlation, and statistical testing using a specially constructed null distribution. We present representative examples to illustrate the impact of these two methodological choices, with **Supplementary Figure S10** showing a gene-metabolite pair that is significant by PCC (FDR < 0.05) but not by SCI and **Supplementary Figure S11** shows a gene-metabolite pair where the SCI is significant (FDR < 0.05) if tested through standard randomization but not via autocorrelation-preserving randomization (FDR > 0.1). Visual inspection of the feature maps reveals no clear spatial covariation in either example. These analyses and examples show that the compute_genemet_sci provides a more conservative test of gene-metabolite spatial covariation compared to the standard correlation approach.

### Applications to human multi-omics data sets

We applied MANTIS to a published human lung-cancer multi-omics dataset [1] that contains 190 metabolites and 22 proteomic markers measured at single-cell resolution across ∼920 cells (**Figure 5A**). We did not have ready access to cell type or regional annotations, so we directly used the “compute_genemet_sci” function, treating protein abundances as a measure of gene expression. This revealed 547 significant protein-metabolite pairs (FDR < 0.05), i.e., ∼13% of 4180 tested pairs (**Figure 5B, Supplementary Table S14**). T-cell markers CD45RA and CD45RO were among the most metabolite-associated proteins, suggesting that immune infiltration and activation states are closely tied to spatial metabolic structure [30,31]. E-cadherin, an epithelial cell marker, is also highly metabolite-associated, potentially reflecting the distinct metabolic programs linked to EMT and/or tumor-stroma boundaries [32,33]. The many associations of Histone H3 may reflect simply a higher metabolite abundance in cell-dense regions or a more specific link to chromatin state. In **Figures 5C,D**, we see that the metabolite glycine (m/z 74.0) has significantly higher expression in tumor regions, defined by ECadherin and PanKeratin positive cells. This is consistent with established mechanisms whereby tumor cells up-regulate glycolysis and one-carbon (serine/glycine) metabolism to support rapid growth, nucleotide and protein synthesis, resulting in elevated glycine accumulation in tumor zones [34]. As a second example, CD45RA (T cell marker) expression positively correlates with ceramide-phosphocholine (m/z 101.961) abundance (**Figures 5E,F**). This may reflect local remodeling of sphingolipids in immune infiltration, consistent with known roles of sphingolipids their ceramide products in T cell activation and maintenance [35].

**Figure 5.**
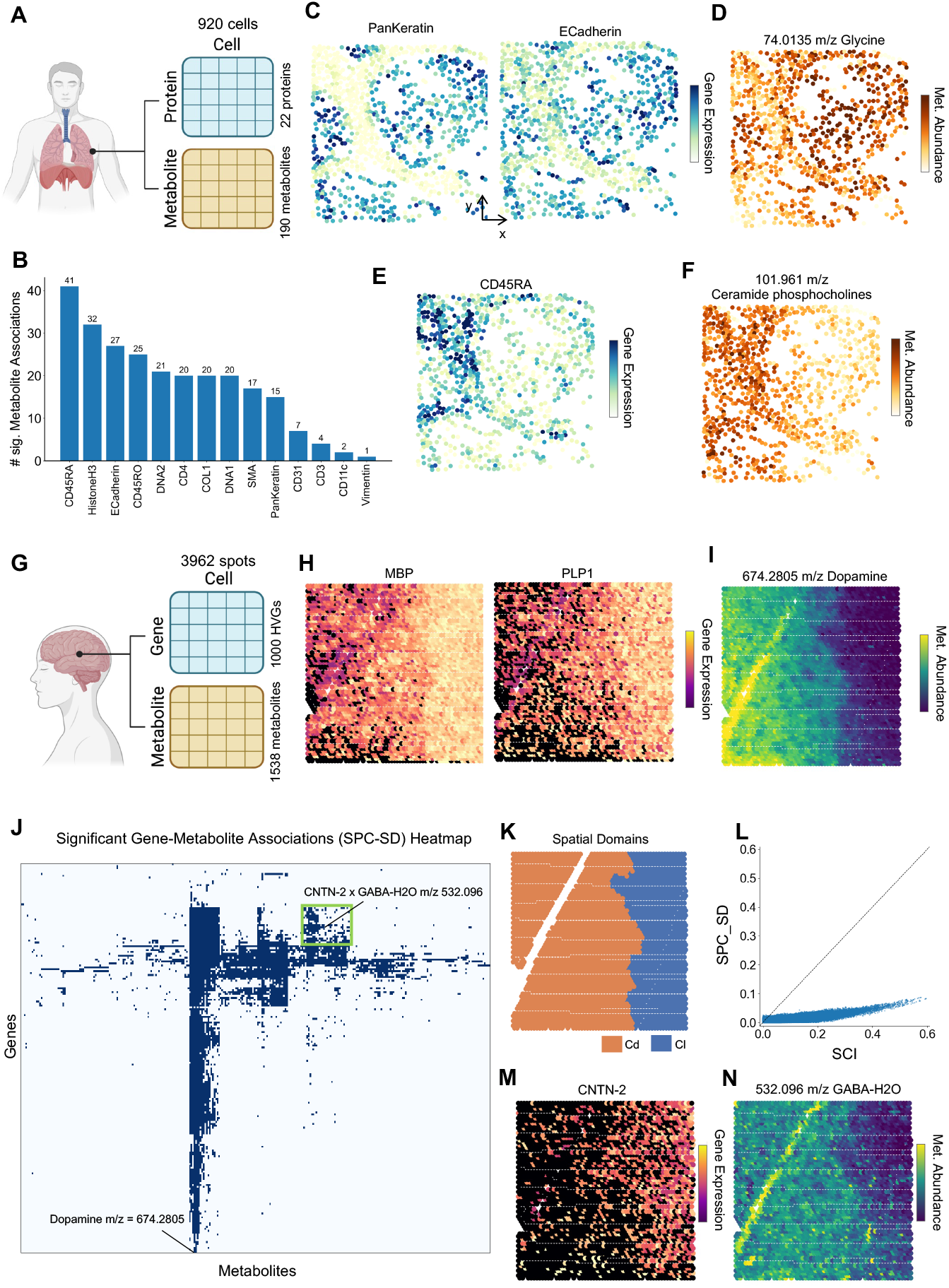
Gene-metabolite spatial covariation in human datasets **(A)** Schematic of the single cell SM + spatial proteomics (SP) human lung cancer sample (920 cells, 22 proteins, 180 metabolites). **(B)** Number of significant protein–metabolite associations identified for each protein. **(C)** Spatial expression patterns of the tumor-associated proteins PanKeratin and E-Cadherin. **(D)** Spatial abundance of glycine (m/z 74.0135), which colocalizes with epithelial tumor marker proteins. **(E)** Spatial expression of the immune T-cell marker CD45RA. **(F)** Spatial abundance of ceramide phosphocholine (m/z 101.961), which shows strong correspondence with CD45RA. **(G)** Schematic of the ST+SM human striatum sample (3962 spots, 1000 HVGs, 1538 metabolites). **(H)** Spatial expression of oligodendrocyte-associated genes *Mbp* and *Plp1*. **(I)** Spatial distribution of dopamine (m/z 674.2805), which is strongly enriched in regions expressing *Mbp*/*Plp1*. **(J)** Heatmap of significant gene–metabolite associations identified by SPC–SD, highlighting broad associations involving dopamine as well as a more localized association involving *Cntn2* and GABA·H2O (m/z 532.096). The green box marks the bicluster from which the *Cntn2*–m/z 532.096 example was selected. (K) Two spatial domains in this sample. Cd: caudate, Cl: claustrum. **(L)** Comparison of SCI and SPC–SD scores for all gene–metabolite pairs. **(M)** Spatial expression of *Cntn2*. **(N)** Spatial abundance of m/z 532.096, showing colocalization with *Cntn2*.

We next applied MANTIS to a human striatum spatial multi-omics sample [5] of postmortem tissue from a 94-year-old Parkinson’s disease patient, examining associations between the top 1000 highly variable genes and all 1538 detected metabolites (**Figure 5G**). The “compute_genemet_sci” function revealed extensive gene-metabolite associations, with 25% of the tested pairs being significant at FDR < 0.05 (**Supplementary Table S15**). Many of these associations involved the metabolite dopamine (m/z 674.281, **Figure 5I**), including a strong negative spatial correlation with *Mbp* (Myelin Basic Protein) and *Plp1* (Proteolipid Protein 1) (**Figure 5H**), which are major myelin-sheath proteins produced by oligodendrocytes and are enriched in white matter but largely absent in grey matter. Regions with high dopamine signal correspond to the caudate nucleus, a gray-matter region rich in neurons, whereas regions with low dopamine signal show white-matter characteristics with higher *Mbp* and *Plp1* expression. This spatial segregation likely explains the observed negative correlation, which did not remain significant (SCI = 0.08) after regressing out the spatial domain.

To test whether other reported associations were driven by spatial domains, we used the “compute_SPC-SD” function that accounts for spatial domains Caudate nucleus and Claustrum (**Figure 5K**). The SCI values fell sharply (**Figure 5L**) and only 5,079 (0.3%) of gene–metabolite pairs remained significant (FDR < 0.05) (**Supplementary Table S16**), indicating that most of the originally observed co-localizations are attributable to spatial domains. **Figure 5J** provides an overview of the gene-metabolite pairs that remain significant, showing that dopamine (m/z = 674.2805) has among the most number of associations even after accounting for domain effects. We noticed a few dense biclusters (modules) of gene–metabolite pairs in the map, e.g., one shown by green box in Fig. 5J. **Figures 5M,N** visualize a representative pair from this module (*Cntn2* (Contactin-2) paired with GABA-H2O (m/z 353.1652), FDR = 0.031), showing a significantly negative spatial correlation in the top right rather than following the spatial domain pattern of Fig. 5K. *Cntn2* is associated with myelinated axons [36] and oligodendroglial structures, whereas GABA-H2O reflects GABAergic neuronal activity [37]. Because fiber-enriched, oligodendroglial compartments and GABAergic neuron–rich compartments rarely coincide in the same tissue locations, *Cntn2* and GABA-H2O exhibit an inverse spatial relationship.

## DISCUSSION

There is rapidly growing interest in tools for analysis of spatial metabolomics (SM) data [2-4], especially those that analyze matched SM and ST data on the same biological sample [1,8,11-13,38]. MANTIS is a computational toolkit for analysis of such co-registered SM+ST data sets at cell/spot resolution, along with cell type annotations and regional delineations, if available. Its main functionalities include identifying metabolites with statistically interesting spatial patterns and discovering gene-metabolite relationships via correlation analysis.

MANTIS adopts new statistical approaches designed to handle confounders and control false positive errors. For example, it uses spatial cross-correlation to discover gene-metabolite relationships, in contrast to standard correlation measures used by SpaMTP and SpatialMETA. Prior work makes a compelling case for use of spatial statistics when quantifying associations in spatial data, omics [39] and otherwise [40], and our empirical comparisons show that a large number of associations reported by non-spatial analysis are not significant when using spatial scores (Supplementary Table S17). Furthermore, MANTIS’ special method for significance testing, based on autocorrelation-preserving permutations, distinguishes it from simple correlation measures implemented in existing tools, and provides a stringent control on false positive errors.

Another important methodological contribution of MANTIS is its handling of potential confounders when testing associations. A gene and metabolite may be spatially co-expressed because they are individually associated with the same spatial region or cell types. Our decision was to report such associations separately from those pairs whose spatial covariance cannot be explained by simple regional or cell type associations of either molecule. Thus, the “compute_SPC-CT” function reports significant gene-metabolite pairs after removing the effect of spatial domains or cell types on a pair’s association. Similar handling of confounders is also featured in MANTIS the function “spatvar_met” for discovering “spatially variable metabolites” (SVM). This function may be employed with varying length scale parameter *l* to reveal patterns at different spatial scales.

The goals and functionalities of MANTIS are different from an emerging genre of tools such as MISO [1,13,41] that combine single-cell metabolomics and gene expression data to obtain a unified multi-omic representation of each cell, which is then used to discover spatial neighborhoods with characteristic molecular signatures. However, these tools do not directly address the explicit recovery of gene-metabolite relationships in a spatial omics setting.

There are several methodological aspects of the MANTIS toolkit that need future research and improvement. For instance, the SCI score is better suited for spot-level data where each spot is assigned cell type proportions (real valued), but may not be ideal when cell type annotations are categorical or binary. For such situations, a spatial covariance test tailored for discrete variables, such as the Proximal Pairs test [42], may be more appropriate. Another direction of future work will be to improve how spatial autocorrelation is accounted for when quantifying bivariate spatial correlation. Currently this is addressed by estimating an empirical null distribution of the SCI score that conditions on metabolite autocorrelation levels, but future versions may explore a modified version of the SCI score that is normalized by the autocorrelation levels of both variables.

MANTIS is intended to add to the developing ecosystem of tools for SM+ST data analysis. These tools offer several useful functionalities, including multi-omics data alignment and mapping, clustering and discovery of markers, and pathway analysis. Our goal was to improve the rigor and precision of spatial pattern discovery and association analysis, and we believe MANTIS presents an important step forward in this direction.

## METHODS

### Data source and preprocessing

We used the dataset from Vicari et al. [5], which provides matched ST (Visium) and SM (MALDI-MSI) data from mouse and human striatum. Gene expression and metabolite abundance measurements are already aligned to the same coordinate system, at the resolution of Visium spots. Also included are annotations of spatial domains and cell type proportions (obtained using Stereoscope [15]) at each spot. We analyzed one mouse striatum tissue section (V11L12-038), with data on 19,126 genes and 2,754 metabolites across 2,493 spatial spots. Raw transcript counts were normalized by library size and log-transformed. We applied MAGIC [43] to reduce technical noise. For each gene, observations outside the interval [*Q*_1_ − 1.5 IQR, *Q*_3_ + 1.5 IQR] were classified as outliers and corrected by local averaging. For metabolomics data, intensities were median-normalized, log-transformed, and corrected for outliers in the same way. Then, we selected the top 1,000 highly variable genes and 500 highly variable metabolites (HVMs) for downstream analysis. The human sample (postmortem striatal tissue from a Parkinson’s disease patient) includes 3,962 spots, 14,826 genes and 1,538 metabolites. Preprocessing was as above, and we selected 1000 highly variable genes (HVGs) and all 1538 metabolites for analysis. We also analyzed a lung cancer dataset from Hu et al. [1], where levels of 190 metabolites and 22 proteins (cell type markers) are measured in 920 cells. We used the gene–expression matrix accompanying the publication without further normalization.

### Regional (spatial domain) enrichment analysis: regional_met function

#### KL divergence

To identify metabolites enriched in specific spatial domains (SD), we constructed regional abundance profiles for each metabolite. For metabolite *m*, we first computed its mean abundance in region *r*, and normalized these across regions to form a probability vector {*p*_*rm*_}_*r*_. A background profile *q*_*r*_ was similarly defined by averaging across all metabolites. We then quantified the divergence between each metabolite’s profile *p*_*m*_ and the background profile *q* using Kullback–Leibler (KL) divergence: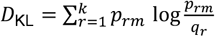, and the contribution of each region was given by the term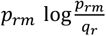.

#### Significance of KL divergence score

To construct the null model, we generate one “null spatial map” 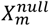 for each metabolite *m* – a randomized version of the metabolite’s spatial distribution where the marginal distribution as well as spatial autocorrelation are preserved – using the sampling procedure described below. We compute the KL divergence score based on 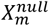 as for the real data. Collecting these scores for all metabolites we obtain an empirical null distribution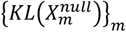. We then fit an analytical distribution to this empirical null, through maximum likelihood. For this, we screen a panel of candidate distribution families, fitted using their standard parameterizations as implemented in scipy.stats and choose the best fit. We evaluate significance for each metabolite *m* by comparing its observed *KL* divergence to the fitted CDF and obtaining one-sided p-value, which is then corrected for multiple testing across metabolites with Benjamini– Hochberg FDR procedure.

### Simulated-annealing (SA) sampler of null spatial maps

We generate null spatial map of a metabolite by repeated swaps of its abundance between pairs of objects (spots/cells), which preserves the marginal distribution. The procedure strives to retain the autocorrelation level of the original spatial map, as follows. A weighted spatial graph *G* is first built from object coordinates, with adjacent objects being connected by edges. Let *i* and *j* index two nodes (objects) in the graph, we write (*i, j*) ∈ *E* if the two spots are neighbors (Euclidean distance *d*_*ij*_ < *l*), *l* being a user-specified length scale with default value *l* = 4*d*_*min*_, where *d*_*min*_ is the shortest distance between spots/cells. Edge (*i, j*) is assigned a weight *w*_*ij*_ as follows:

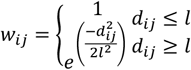

Let *X*_*m*_ = {*X*_*im*_}_*i*_ denote the spatial map of a metabolite *m* across all spots *i*. Let *M*(*X*) be the autocorrelation of any map *X*, i.e., *M*(*X*) = ∑_(*i,j*)∈*E*_ *w*_*ij*_ *X*_*i*_ *X*_*j*_. The sampler explores randomized maps 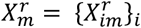 such that the autocorrelation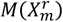 remains close to the autocorrelation *M*_0_ = *M*(*X*_*m*_) of the original map *X*_*m*_. In each iteration, the sampler evaluates a proposed swap of metabolite levels between two randomly chosen nodes *i, j*. Denoting the current spatial map by *X*^*curr*^ and the autocorrelation of the update map after the swap by *M*_*new*_, the proposal is accepted with probability *α* = min[ 1, exp(−(‖*M*_*new*_ − *M*_0_‖_1_ − ‖*M*(*X*^*curr*^) − *M*_0_‖_1_)/*T*)], where *T* is the current “temperature” of the SA sampler.

See Supplementary Methods S1 for additional details of implementation of the SA sampler of randomly permuted spatial maps.

#### Outputs

Each SA run returns the final permutation *π* of node indices obtained by the sequence of swaps performed.

### Spatial Cross-Correlation Index (SCI)

#### Definition

For vectors ***x, y*** ∈ *R*^*N*^ and a spatial weight matrix *W* = [*w*_*ij*_], we define the Spatial Cross-Correlation Index (SCI) as a bivariate Moran-type correlation: 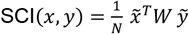, where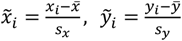, and *N* is the number of objects (spots/cells). Here,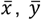 denote sample means and *s*_*x*_, *s*_*y*_ are sample standard deviations.

#### Uses of SCI

In all uses of SCI, *y* is the metabolite intensity. For celltype_met, *x* is cell-type proportion (for spot-based data) or a binary indicator of cell type (for single-cell data). For “compute_genemet_sci”, *x* is gene expression.

*Weight matrix W* is defined as:

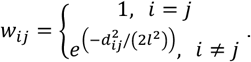

We row-normalize the matrix, i.e., ∑_*j*_ *w*_*ij*_ = 1. The default value of *l* is *d*_*min*_.

#### Significance testing

We use sampler-based empirical nulls (described above) to control for metabolite spatial autocorrelation when assessing SCI significance. (See **Supplementary Methods S2**.)

### Detection of spatially variable metabolites (SVM)

#### Moran’s I

We quantify spatial autocorrelation of a metabolite’s distribution {*X*_*im*_}_*i*_ using a revised Moran’s I. Let W= [*w*_*ij*_] with *w*_*ii*_ = 0 and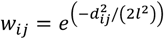for *i* ≠ *j*. We row-normalize W such that 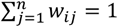 for all i. Under this row-normalized formulation, Moran’s I for metabolite *m* is given by 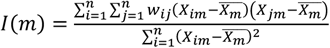.

#### Control for cell type associations

We first regress the metabolite abundance against cell type. Specifically, we fit a LASSO model to {*X*_*im*_}_*i*_ using cell type(s) of object *i* as covariates and select a sparse subset of cell types informative of *X*_*im*_. Then we refit an OLS model on the selected cell type covariates and calculate Moran’s I as above on the residuals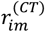 of the metabolite, to obtain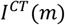.

#### Control for spatial domain association

We encode each obje *f f f* ct’s region (spatial domain) annotation with a one-hot feature vector, which is used to predict metabolite abundance via an OLS model and obtain residuals 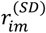, on which we then calculate Moran’s I as above to obtain *I*^*SD*^(*m*).

We designate a metabolite as SVM if min(*I*^*CT*^(*m*), *I*^*SD*^(*m*)) ≥ *τ* for some threshold *τ*.

### Spatial Partial Correlation (SCI with adjustment for cell type or spatial domain)

#### Adjustment for cell type/spatial domain

For both the gene expression matrix and metabolite abundance matrix, we apply the same adjustments as above: spatial domains by OLS and cell types by LASSO with post-selection OLS refit. This yields residuals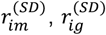of metabolite abundance and gene expression respectively after accounting for effects of spatial domains, and similarly the residuals 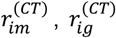 after accounting for cell types. We define “spatial partial correlation” (SPC) as the SCI score between residuals, i.e. 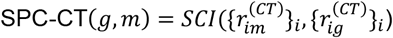 and 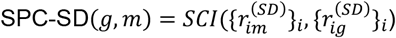

#### Null model and statistical testing of SPC

We use the SA sampler to obtain a randomized version *X*^*r*^ of each metabolite’s spatial map *X*, and the full collection of randomized spatial maps is then subjected to SPC-SD or SPC-CT calculations, thus resulting in null distributions and empirical p-value (as well as BH-FDR q-value) calculations as above.

## KEY POINTS

- MANTIS is an analytics toolkit for paired spatial metabolomics + spatial transcriptomics data, supporting discovery of metabolic spatial patterns and gene-metabolite relationships.
- MANTIS quantifies gene-metabolite spatial associations using a spatially aware correlation score and performs autocorrelation-preserving permutation testing to provide rigorous statistical significance and reduce false positives.
- MANTIS disentangles confounding effects of spatial domains and cell-type composition using spatial partial correlation, separating domain/cell-type-driven co-localization from more specific associations.

## ACKNOWLEDGMENTS

We thank Dr. Ahmet Coskun and Efe Ozturk for helpful discussions and sharing data.

## DATA AVAILABILITY

The datasets used in this study were obtained from publicly available sources and are listed in Supplementary Table S18. MANTIS is implemented in Python and distributed as open-source software at https://github.com/yuhaotuo/MANTIS

## Notes

### Competing Interest Statement

The authors have declared no competing interest.

### Summary of Updates

This version includes revisions to the abstract to improve clarity and better summarize the study objectives, methods, and key findings. The acknowledgments section has been updated. The supplementary tables have been revised and updated to reflect these changes. No substantial changes have been made to the main text, figures, or conclusions.

### Funding

- National Institutes of Health
- U.S. National Science Foundation

