## Supplementary Information for "MANTIS: Analytics toolkit for spatial metabolomics with matching spatial transcriptomics data"

**Supplementary Table S1. Region-associated metabolites detection in the VICARI data set**. For each metabolite (reported by m/z), we report the KL-divergence of its regional abundance profile and the significance of enrichment computed against a spatially aware null. The null for each metabolite is generated using a simulated annealing sampler that preserves the metabolite's global spatial autocorrelation while randomizing its spatial arrangement. Empirical $p$-values ('p_val') are converted to $q$-values with Benjamini-Hochberg FDR control ('q_val'). Metabolites with q < 0.1 were considered region-associate, and only significant associations are listed.

**Supplementary Table S2. Region-associated metabolites identified by Wilcoxon tests across brain regions in the VICARI data set.**

This table provides a baseline comparison to MANTIS by listing region-associated metabolites that are detected as significant in a two-sided Wilcoxon test. For each region, a two-sided Wilcoxon rank-sum test was performed to compare metabolites intensities in the target region versus all other regions. Each row corresponds to a region–metabolite pair. Reported values include the raw p-value ('pval'), the $log_2$ fold change of median intensities between the target region and all others ('$log_2FC$') and the Benjamini-Hochberg adjusted q-value ('qval'). Metabolites with q < 0.05 were considered region associated, and only significant associations are listed.

**Supplementary Table S3. Cell type-associated metabolite detection in the VICARI dataset.**

The table presents metabolites whose spatial patterns show significant correlation with specific cell types. For each cell type-metabolite pair, the Spatial Correlation Index ('SCI') quantifies the spatial co-localization between the metabolite intensity map and the cell-type spatial probability map. Associated p-values ('pval') were obtained from a simulation-based null model, and q-values ('qval') denote FDR-adjusted significance levels. Associations with q < 0.05 were considered significant, and only significant pairs are listed.

**Supplementary Table S4. Baseline spatial association between cell type and metabolite intensities in the VICARI dataset.**

This table reports baseline spatial associations between cell types and metabolites quantified using the Spatial Correlation Index (SCI) in the VICARI dataset. Statistical significance was assessed using a non-spatial shuffled null model, in which values were randomly permuted without preserving spatial structure. For each cell type–metabolite pair, the SCI score quantifies the degree of correspondence between the cell-type probability map and the metabolite intensity map. P-values ('pval') were computed under the shuffled-null distribution of SCI, and q-values ('qval') represent FDR-adjusted significance. Pairs with q-value < 0.05 were considered significant and are listed here.

**Supplementary Table S5. Spatially variable metabolite identification in the VICARI dataset.**

This table presents the identification of metabolites whose spatial autocorrelation remains high even after adjusting for associations with spatial domains and cell types. For each metabolite, we report the baseline Moran's I value (Moran's I) and the adjusted Moran's I values after regressing out spatial domains (I_after_region) and cell types (I_after_celltype). The minimum of the adjusted scores (I_after_min) represents the remaining spatial autocorrelation after correction, and the corresponding reduction from the baseline value (Difference) is also reported. Binary flags (before_flag, region_flag, celltype_flag) indicate whether each Moran's I value exceeds the significance threshold (> 0.2), and the "identification" column marks metabolites designated as spatially variable Metabolite (SVM). Metabolites with minimum of the adjusted scores (I_after_min) > 0.2 were considered SVMs, and only those identified as SVMs are listed.

**Supplementary Table S6. Effect of the spatial length-scale ($l$) on SVM detection rates in the VICARI data set.**

This table presents the impact of varying the Gaussian kernel length-scale on the identification of spatially variable metabolites. For each choice of length-scale, expressed in multiples of the minimum inter-spot distance (dmin), we report the total number of metabolites analyzed (n_metabolites) and the number ('count_>$\tau$') and fraction ('frac_>$\tau$') whose Moran's I values exceed thresholds ($\tau \in \{0.15, 0.20, 0.30\}$). These results show how the choice of $l$ influences the proportion of metabolites identified as spatially correlated.

**Supplementary Table S7. SpatialDE analysis of metabolite spatial variability in the VICARI data set.**

This table presents the SpatialDE results for each metabolite across spatial spots. For each metabolite, the fraction of spatial variance ('FSV') estimates the proportion of total variance explained by a spatial covariance component, where higher FSV indicates a larger spatial component of variation. The length-scale parameter ($l$) quantifies the spatial correlation range of the fitted Gaussian-process model, and the variance term (s2_FSV) is representing the portion of gene-expression variability not explained by spatial structure. Model statistics include the log-likelihood ratio ('LLR') and significance is assessed via p-value ('pval') and FDR-adjusted q-value ('qval'). Metabolites with q < 0.05 were considered significant, and all significant associations identified by SpatialDE are listed.

**Supplementary Table S8. Gene-metabolite spatial correlations without corrections in the VICARI data set.**

This table presents significant gene-metabolite pairs (q < 0.05) whose Spatial Correlation Index (SCI) indicates spatial co-localization between the gene expression and the metabolite abundance without applying region or cell-type corrections. For each pair, the SCI score (SCI) quantifies spatial co-expression across spots, while the metabolite bin used in testing is noted (bin_index), and each metabolite is partitioned into ten equal-frequency bins based on its spatial autocorrelation. The empirical p-values (p_value) are derived from a simulation-based null that preserves metabolite spatial autocorrelation, and the FDR-adjusted q-values (q_value) indicate statistical significance. Pairs with q < 0.05 are marked as significant (significant), and only significant associations are listed.

**Supplementary Table S9. Cell type-corrected gene-metabolite spatial correlations in the VICARI data set.**

This table presents significant gene-metabolite pairs identified after regressing out cell type effects and computing the Spatial Partial Correlation. For each pair, 'SPC_CT' quantifies the spatial association remaining after removing cell type-driven confounding effect, while empirical p-values (pval) are derived from a simulation-based null that preserves metabolite spatial autocorrelation, and q-values (qval) represent FDR-adjusted significance. The bin used for testing is recorded (bin_id), and pairs with q < 0.05 are marked as significant (significant).

**Supplementary Table S10. Gene-metabolite association explained by cell-type effects.**

This table presents gene-metabolite pairs that are significant in the baseline Spatial Correlation Index (SCI) analysis but lose significant after cell-type correction (SPC_CT), indicating that their spatial co-localization is primarily driven by underlying cell-type composition rather than independent spatial coupling. For each pair, 'SCI' and 'SPC_CT' report the spatial correlation before and after regressing out cell-type effects, and empirical significance is assessed through p-values and q-values for both analyses (p_value_SCI, q_value_SCI, pval_SPC_CT, qval_SPC_CT). The columns 'significant_SCI' denote significance at q < 0.05, and 'significant_SPC_CT' denote significance at q < 0.1; only pairs significant under SCI but not under SPC_CT are included.

**Supplementary Table S11. Domain-corrected gene-metabolite spatial correlations after regressing out spatial-domain effects in the VICARI data set.**

This table presents significant gene-metabolite pairs identified after regressing out spatial-domain effects and computing the Spatial Partial Correlation. For each pair, **'**SPC_SD' quantifies the spatial association remaining after removing domain-driven covariance, while empirical significance is assessed through p-values ('pval') derived from a simulation-based null and FDR-adjusted q-values ('qval'). The bin used for testing is recorded ('bin_id'), and gene–metabolite pairs with q < 0.05 are marked as significant ('significance'), and only pairs significant pairs are listed.

**Supplementary Table S12. Gene-metabolite spatial correlations independent of cell-type and spatial domain effects in the VICARI data set.**

This table presents gene-metabolite pairs that are jointly significant in both the cell type-corrected analysis (SPC_CT) and the domain-corrected analysis (SPC_SD). For each pair, column 'SPC_CT' and 'SPC_SD' report the spatial correlation scores after adjusting for cell-type and domain effects, respectively, and empirical significance is assessed through corresponding p-values and q-values ('pval_CT', 'qval_CT', 'pval_SD', 'qval_SD'). Significance indicators denote pairs with q < 0.05, and the overlap column marks pairs significant in both analyses. Only gene–metabolite pairs passing both criteria are listed.

**Supplementary Table S13. Cumulative number of significant gene–metabolite pairs across q-value thresholds based on the baseline Pearson correlation analysis.**

This table presents baseline, non-spatial associations between genes and metabolites using the Pearson correlation coefficient. It summarizes the cumulative number of significant gene–metabolite pairs at varying q-value thresholds. For each threshold, the count represents the total number of pairs with FDR-adjusted q-values below the specified cutoff. These q-values were derived from Pearson correlation analyses assessing baseline, non-spatial associations between genes and metabolites across spatial spots.

**Supplementary Table S14. Protein-metabolite spatial correlations without corrections in the human lung cancer data set.**

This table presents significant gene-metabolite pairs from the human lung cancer dataset. Using the Spatial Correlation Index ('SCI') computed without cell type or domain correction, spatial co-localization is quantified between the protein-abundance and the metabolite-abundance. The bin used for testing is recorded (bin_id), empirical p-values ('pval') are obtained from a simulation-based null that preserves metabolite spatial autocorrelation, and FDR-adjusted q-values ('qval') denote significance. Protein-metabolite pairs with q < 0.05 are marked as significant ('significant'), and only significant associations are listed.

**Supplementary Table S15. Gene-metabolite spatial correlations without corrections in the human striatum data set.**

This table presents significant gene–metabolite pairs from the human striatum dataset based on the Spatial Correlation Index (SCI) computed without cell type or domain correction. For each pair, column 'SCI' quantifies spatial co-localization between the gene-expression map and the metabolite-abundance map within the human striatal sections. The bin used for testing is recorded ('bin_id'), empirical p-values ('pval') are computed within each bin using a simulation-based null, and FDR-adjusted q-values ('qval') denote significance. Gene-metabolite pairs with q < 0.05 are marked as significant ('significant'), and only significant associations are listed.

**Supplementary Table S16. Domain-corrected gene-metabolite spatial correlations after domain removal in the human striatum data set.**

This table presents significant gene–metabolite pairs from the human striatum dataset based on based on the domain-corrected Spatial Partial Correlation (SPC_SD). For each pair, 'SPC_SD' quantifies spatial co-localization between the gene-expression map and the metabolite-abundance map after regressing out spatial-domain effects. The bin used for testing is recorded ('bin_id'), empirical p-values ('pval') are derived from the simulation-based null, and FDR-adjusted q-values ('qval') denote significance. Gene-metabolite pairs with q < 0.05 are marked as significant ('significant'), and only significant associations are listed.

**Supplementary Table S17. Cumulative significant gene–metabolite pairs across q-value thresholds using SCI with random-shuffle null.**

This table reports the cumulative number of significant gene–metabolite associations identified in the human striatum dataset using the Spatial Correlation Index (SCI) at varying q-value thresholds. Statistical significance was evaluated using a random-shuffle null model, and empirical p-values were adjusted for multiple testing using FDR correction. For each threshold, the count reflects the total number of gene–metabolite pairs with q-values below the specified cutoff.

**Supplementary Table S18.**

This table presents the datasets used in this study and their description.

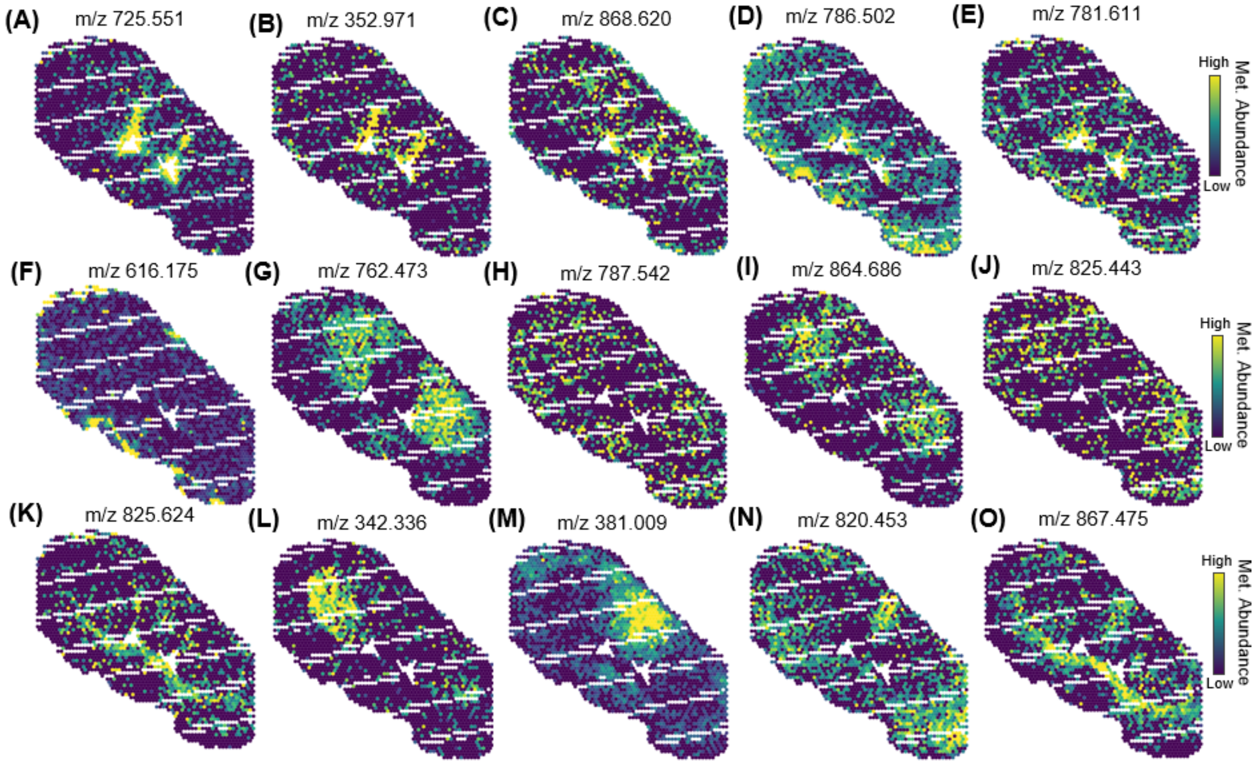


**Supplementary Figure S1**

Metabolite abundance maps for selected region-associated metabolites. Panels **(A–O)** display spatial distributions metabolite intensities (m/z values indicated) across spots. The 15 panels correspond to metabolites summarized in regional-associated heatmap in Figure 2E and visually illustrate the spatial patterns underlying those regional associations.

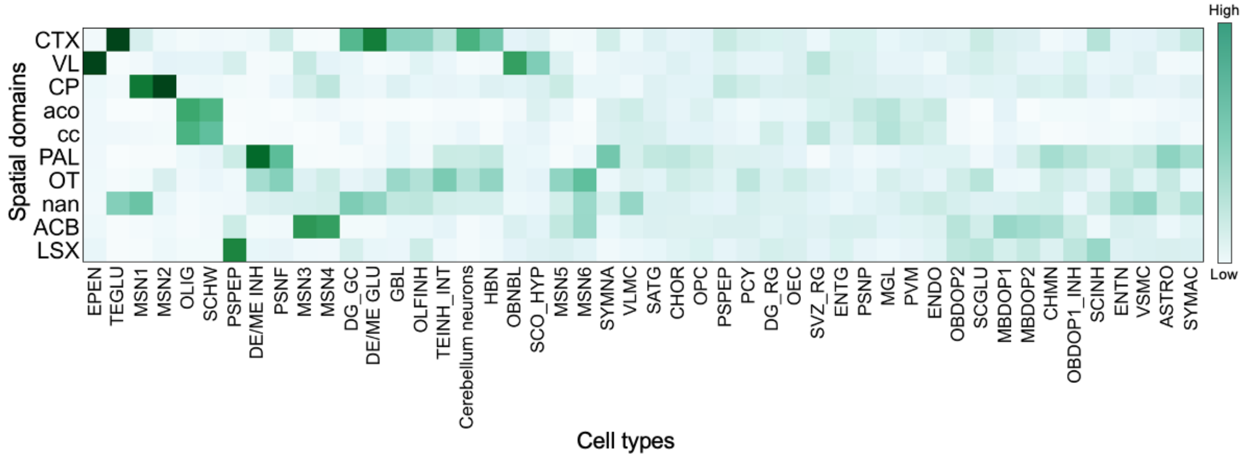


**Supplementary Figure S2**

Clustered heatmap showing regional abundance profiles of cell types. Each row represents one anatomical region, and each column represents one cell type. Colors indicate normalized cell type proportions within each region. Hierarchical clustering (Euclidean distance, average linkage) was used to group cell types with similar regional patterns. The resulting clusters highlight regionally enriched cell types, including telencephalon projecting inhibitory neurons MSN1 in caudate putamen (CP) and ependymal cells in ventral striatum (VL). This figure complements the regional-association heatmap in Figure 2E and supports the observation that certain region-associated metabolites may reflect underlying cell type composition differences across regions.

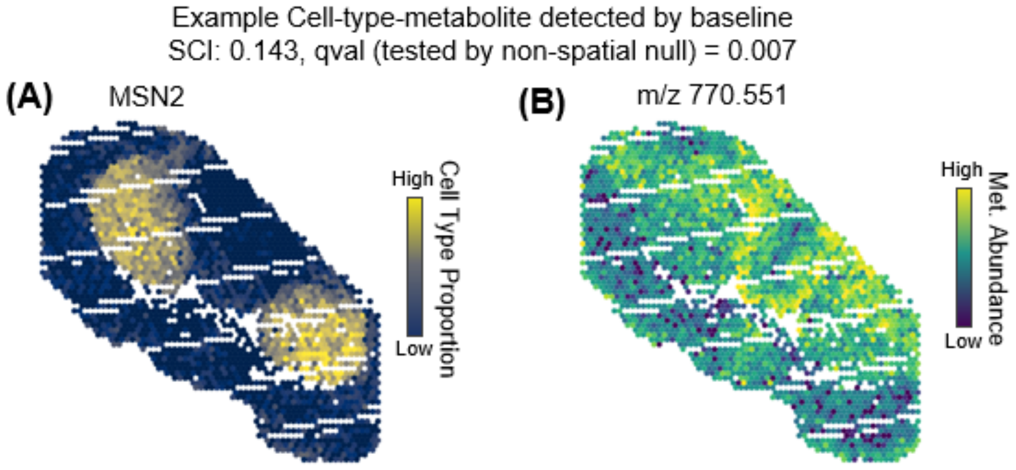


**Supplementary Figure S3**

Example of a cell type-metabolite association identified by a non-spatial baseline (random-shuffled) method (SCI) but not by MANTIS. **(A)** Spatial expression map of MSN2 cell type. **(B)** Spatial abundance map of metabolite m/z 770.551. The baseline SCI test reported a significant association (SCI=0.143, FDR q =0.007). However, the pair was not significant under the MANTIS Spatial Cross-Correlation Index (SCI) test, and visual inspection of the spatial maps does not reveal a clear co-localization between two features. This example illustrates how non-spatial correlation methods may yield false-positive associations when spatial structure is not accounted for.

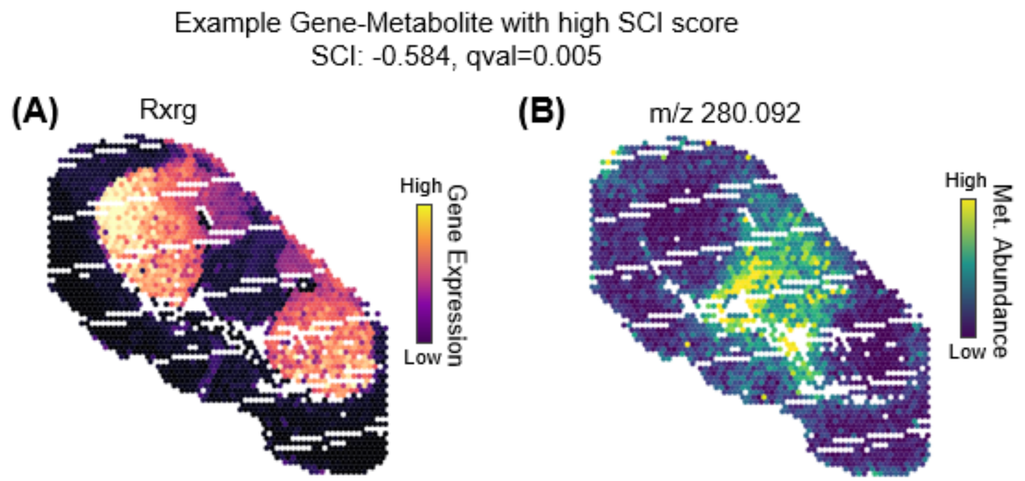


**Supplementary Figure S4**

Example of a gene–metabolite pair with high spatial cross-correlation detected by MANTIS. **(A)** Spatial expression map of *Rxrg*. **(B)** Spatial abundance map of metabolite m/z 280.092. The pair shows a strong negative Spatial Cross-Correlation Index (SCI = −0.584, FDR q = 0.005), indicating clear spatial concordance between gene expression and metabolite abundance. This example illustrates how MANTIS identifies spatially coherent gene–metabolite associations.

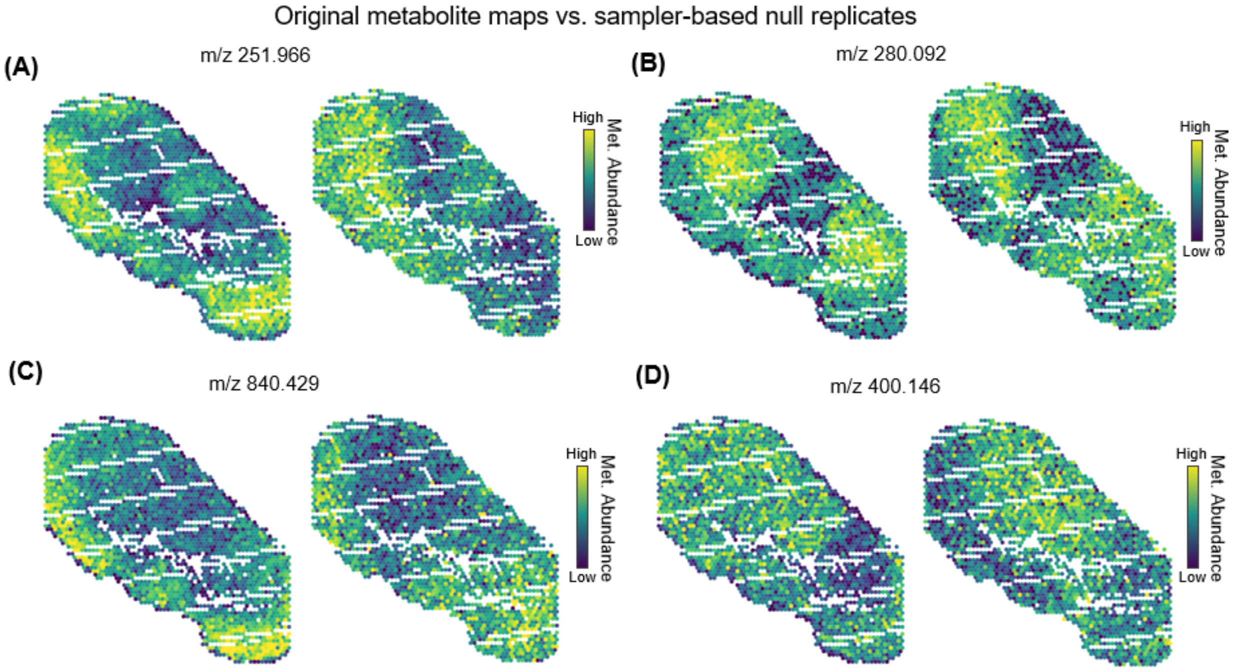


**Supplementary Figure S5**

Comparison of original metabolite maps and sampler-based spatial null replicates. Panels **(A–D)** show four example metabolites used to illustrate the simulated-annealing (SA) sampler procedure. For each example, the left map displays the original metabolite abundance pattern, and the right map shows a single null replicate generated by the MCMC sampler that preserves the metabolite's spatial autocorrelation while randomizing its spatial arrangement. These null replicates maintain the same marginal intensity distribution and neighborhood-level smoothness as the original map but differ in the specific spatial configuration of high- and low-intensity regions, producing realistic yet spatially randomized controls. Color scales are identical within each pair. These null replicates from MCMC sampler collectively form the empirical null distribution used to assess the statistical significance of the Spatial Correlation Index (SCI) in MANTIS.

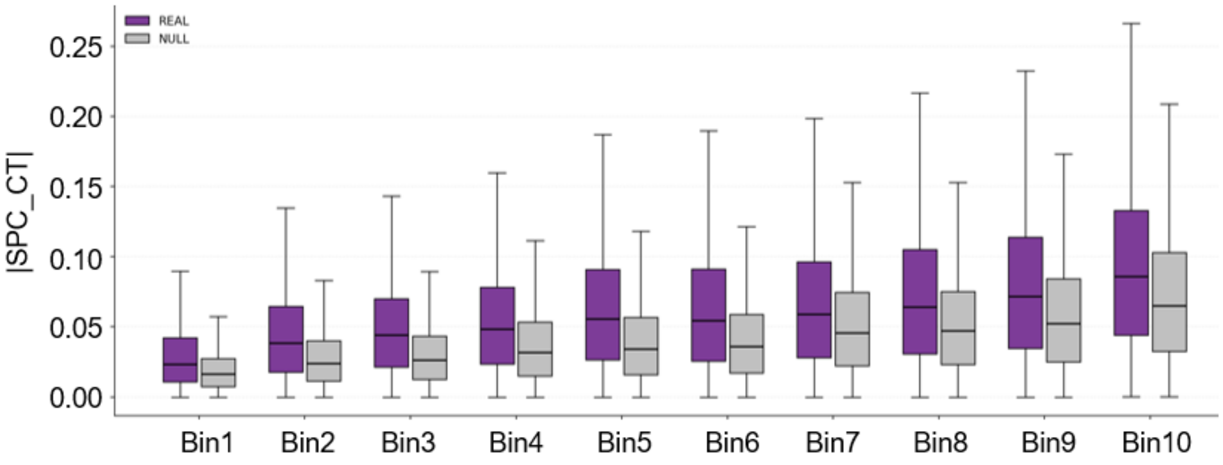


**Supplementary Figure S6**

Comparison of absolute SPC-CT values between real and null distribution across metabolite autocorrelation bins. For each bin representing a range of metabolite spatial autocorrelation (Bin 1 = lowest and Bin10 = highest range), boxplots show the distribution of |SPC-CT| values computed from the real data (purple) and from null replicates generated using the MCMC sampler (gray). Boxes indicate median and interquartile range. The null distribution represents the expected SPC-CT values under randomized spatial organization that preserves spatial autocorrelation but disrupts cell-type–specific coupling. For most bins, many of the real |SPC-CT| values are higher than those from the null distribution, indicating the presence of strong gene-metabolite associations that cannot be explained by cell-type-specific signals (since SPC-CT factors out such signals).

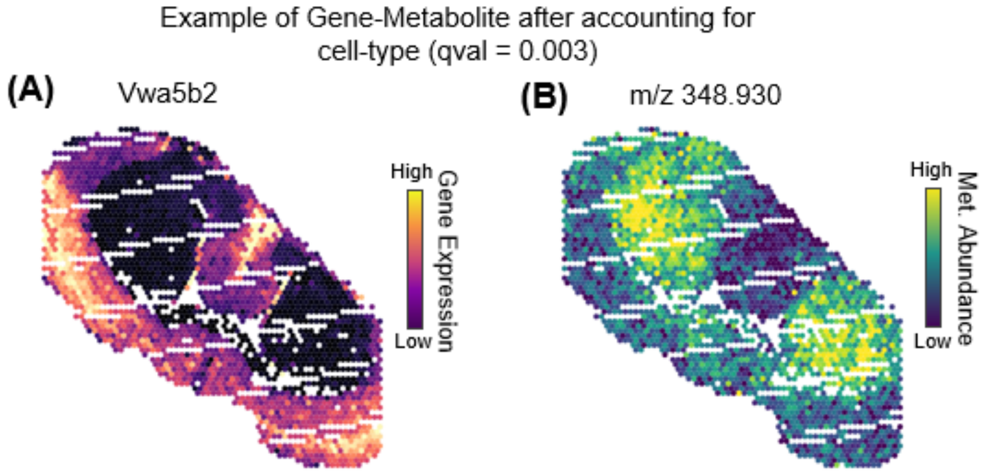


**Supplementary Figure S7**

Example of a gene–metabolite association that remains significant after accounting for cell-type composition. **(A)** Spatial expression map of Vwa5b2. **(B)** Spatial abundance map of metabolite m/z 348.930. After adjusting for cell-type proportions using the “compute_SPC-CT” (Spatial Partial Correlation for Cell Type) function, the association remains significant (FDR q = 0.003). The strong spatial concordance between gene expression and metabolite abundance suggests a cell-type-independent relationship, implying that this correlation is not solely driven by compositional differences among cell types but may reflect an intrinsic spatial or metabolic coupling within the tissue.

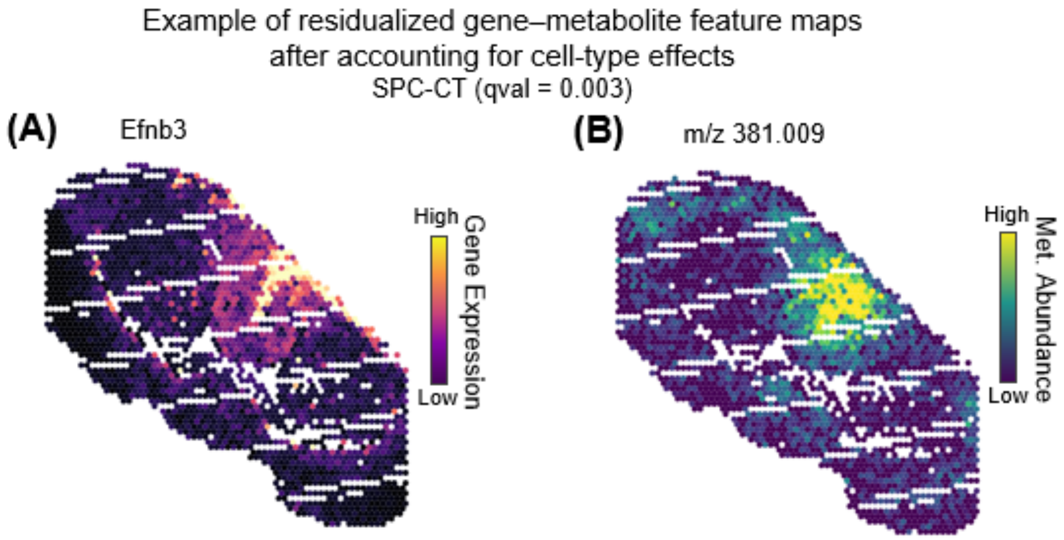


**Supplementary Figure S8**

Residualized feature maps showing a gene-metabolite spatial concordance that persists after regressing out cell-type effects. **(A)** Spatial expression map of Efnb3 residuals (cell-type adjusted). **(B)** Spatial abundance map of metabolite m/z 381.009 residuals (cell-type adjusted). Association remains significant after cell-type regression (“compute_SPC-CT”) (FDR q = 0.003). Both panels show residual values after removing cell-type effects. Significant spatial correlation between the gene and the metabolite remains in the residuals, indicating that the observed co-pattern is not explained solely by cell-type composition but reflects an additional spatially organized relationship within the tissue.

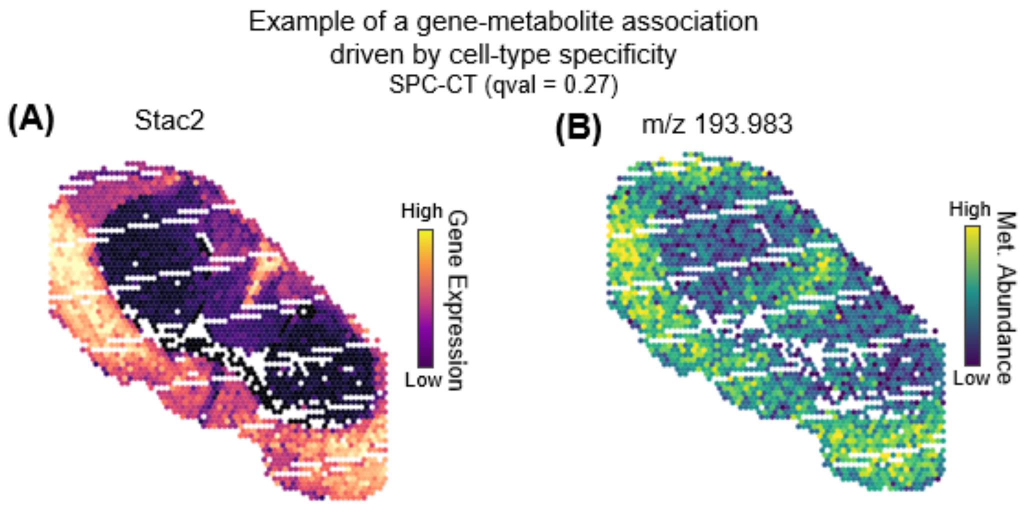


**Supplementary Figure S9**

Example of a gene–metabolite association driven by cell-type specificity (SPC-CT). **(A)** Spatial expression map of Stac2. **(B)** Spatial abundance map of metabolite m/z 193.983. The association shows a highly significant Spatial Correlation Index (SCI) of 0.572 (FDR q = 0.005) before regressing out cell-type effects, but it drops to 0.197 (SPC-CT) (FDR q = 0.27) after adjustment. Although the SCI is large and statistically significant, the lower SPC-CT score and its nonsignificant q-value indicates that the observed association is largely explained by mutual enrichment in a specific cell type (peripheral sensory neurofilament neurons). This example illustrates a cell-type-driven correlation that is removed when controlling for cellular composition using MANTIS.

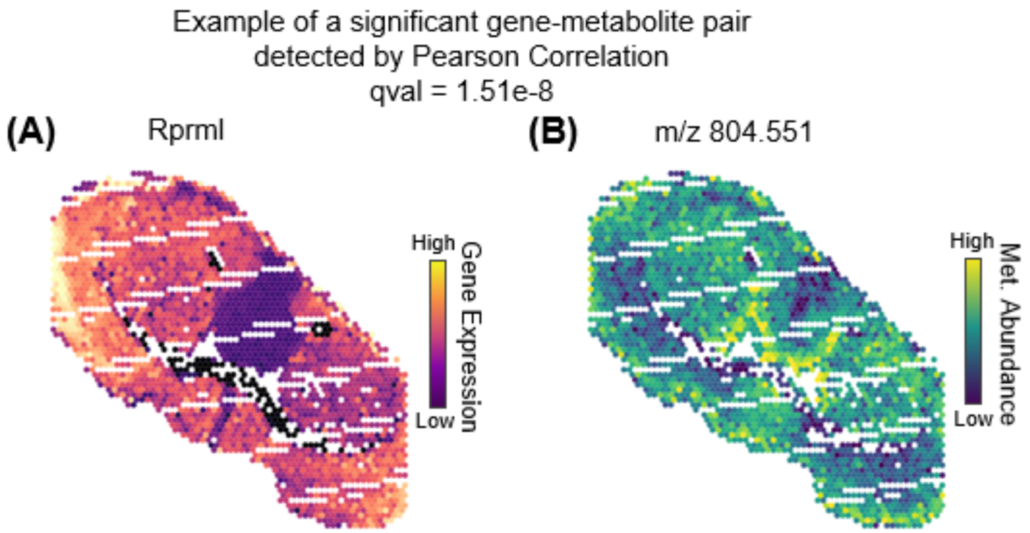


**Supplementary Figure S10**

Example of a gene–metabolite pair that is significant by Pearson correlation but not by spatial correlation (SCI). **(A)** Spatial expression map of *Rprml*. **(B)** Spatial abundance map of metabolite m/z 804.551. The Pearson correlation is highly significant (FDR q = $1.51 \times 10^{-8}$), whereas the Spatial Correlation Index (SCI) (under standard randomization), is not significant (FDR q = 0.109), indicating that the apparent correlation disappears once spatial structure is considered. In panels (A) and (B), although Pearson correlation is strongly significant, this relationship is not visually supported by the spatial feature maps, which show no clear co-localization between gene and metabolite. This example illustrates that a conventional (non-spatial) correlation test can yield false positives in spatial data when spatial structure is ignored.

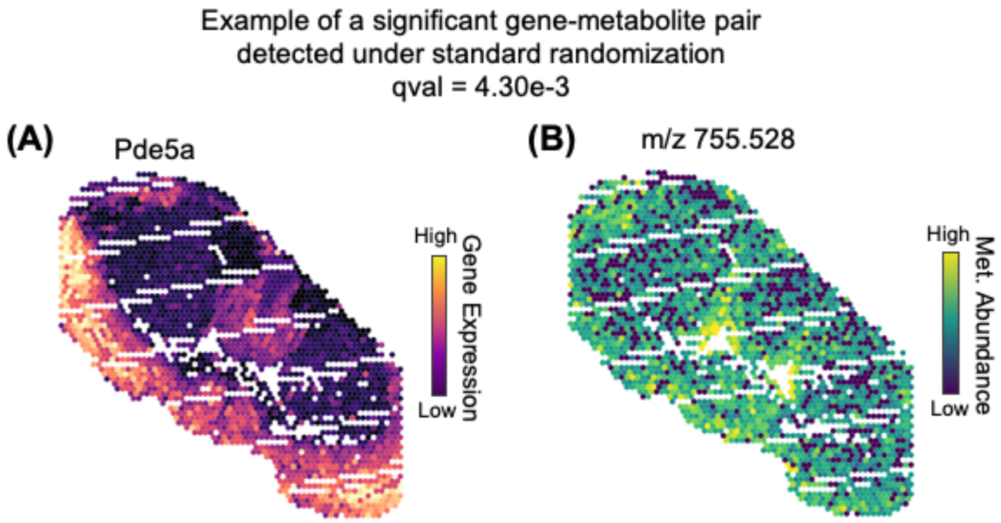


**Supplementary Figure S11.** Example of a gene–metabolite pair that appears significant under standard randomization but not under an autocorrelation-preserving null. **(A)** Spatial expression map of *Pde5a*. **(B)** Spatial abundance map of metabolite m/z 755.528. The Pearson correlation is highly significant (FDR q = $5.01 \times 10^{-8}$), and the SCI is also significant when tested with a standard non-spatial permutation (FDR q = $4.30 \times 10^{-3}$), but it becomes non-significant when evaluated using an autocorrelation-preserving permutation (FDR q = 0.431). Although both PCC and SCI (under standard randomization) appear significant, the spatial maps do not reveal clear spatial concordance. The apparent concordance disappears when spatial structure is properly controlled by autocorrelation-preserving permutation. This example demonstrates that significance detected by non-spatial tests or randomization can be misleading and spatial autocorrelation-preserving randomization is essential to avoid false-positive correlations.

**Supplementary Methods S1. Simulated Annealing (SA) sampler of permuted spatial maps**

We generate null metabolite maps by repeated swaps of metabolite abundance between pairs of objects (spots/cells), which preserves each metabolite's marginal distribution. We adopt a Simulated Annealing strategy to ensure that the randomized map (after many repeats) retains the autocorrelation level of the original.

A weighted spatial graph $G$ is first built from object coordinates, with adjacent objects being connected by edges. Let $i$ and $j$ index two nodes (objects) in the graph, we write $(i,j) \in E$ if the two spots are neighbors, meaning that their Euclidean distance $d_{ij}$ is within a prescribed neighborhood scale $l$. The parameter $l$ is a user-specified length scale that defines the neighborhood radius within which interactions are treated as equally strong, and the default $l = 4d_{min}$. Edge $(i,j)$ is assigned a weight $w_{ij}$ as follows:

$$w_{ij} = \begin{cases} 1 & d_{ij} \leq l \\ e^{\left(\frac{-d_{ij}^2}{2l^2}\right)} & d_{ij} \geq l \end{cases}$$

Let $\{X_{im}\}_i$ denote the spatial map of a metabolite $m$, where $X_{im}$ is the intensity of metabolite $m$ at spot $i$. Let us define $M(X)$ as the autocorrelation of a spatial map $X$, i.e., $M(X) = \sum_{(i,j)\in E} w_{ij}\ X_{im} X_{jm}$. The sampler explores randomized maps $X^r = \{X_{im}^r\}_i$ obtained by repeated swaps of metabolite level between randomly chosen pairs of nodes, such that the autocorrelation $M(X^r)$ remains close to the autocorrelation $M_0 = M(X)$ of the original spatial map $\{X_{im}\}_i$. In each iteration, the sampler evaluates a proposed swap of metabolite levels between two randomly chosen nodes $i, j$. The effect of such a swap is calculated efficiently as follows. Let $X^{curr}$be the current spatial map. Define $S[i,m] = \sum_{k\in Nbr(i)} w_{ik} X_{km}^{curr}$. Then the change in $M(X^{curr})$ due to swap of $i, j$ is given by $\Delta M = \left(X_{jm}^{curr} - X_{im}^{curr}\right)[\left(S[i,m] - w_{ij}X_{jm}^{curr}\right) - \left(S[j,m] - w_{ji}X_{im}^{curr}\right)]$. Let the new autocorrelation if this swap is to be accepted be denoted by $M_{new} = M(X^{curr}) + \Delta M$. The proposal is accepted with probability $\alpha = \min[1, \exp(-(\|M_{new} - M_0\|_1 - \|M(X^{curr}) - M_0\|_1)/T)]$, where $T$ is the current "temperature" of the SA sampler. After an accepted swap, $S$ is updated locally.

*Burn-in:* We perform a pre-shuffle phase of uniformly sampled swaps to decorrelate the initial state and to estimate a scale parameter used in the annealing schedule.

*Annealing schedule & acceptance*: Let $t$ be the base scale (by default set to the mean change in autocorrelation observed during burn-in swaps). The temperature starts at $T_{\text{start}} = t \cdot t\_scale\_start$ and decreases linearly every $t\_scale\_every$ steps to a floor $T_{min} = t \cdot t\_scale\_min$.

*Early stopping*: The sampler uses multiple early-stopping criteria: the energy $E(M) = \|M - M^*\|$ remains below absolute/relative thresholds or does not improve for an extended stretch of accepted swaps, or the total number of proposed swaps exceeds a threshold.

*Outputs*: Each SA run returns the final permutation $\pi$ of node indices obtained by the sequence of swaps performed.

**Supplementary Methods S2: Significance testing of SCI scores**

We use sampler-based empirical nulls to analyze while controlling for metabolite spatial autocorrelation. **Cell type–metabolite SCI.** We fix $x$ (cell-type proportion) and generate a null map for each metabolite by simulated-annealing (SA) permutations of $y$. For each cell type c, we build a cell-type–specific distribution by observed SCI values across all metabolites, and use this distribution to obtain two-sided empirical p-values, which are converted to obtain q-values by the Benjamini–Hochberg procedure. **Gene–metabolite SCI.** We divide the complete collection of metabolites into groups based on similar levels of spatial autocorrelation and construct null distributions of gene-metabolite SCI scores for each such group separately, thus controlling for the effect of metabolite autocorrelation. Let $M_0(m)$ denote the spatial autocorrelation measure of metabolite $m$. We divided the observed range of $M_0(m)$ over all metabolites into 10 equal-frequency "bins". For a metabolite $m$, the observed SCI is compared to the bin-matched null distribution formed by pooling gene-metabolite SCI values from SA-permuted maps of metabolites whose $M_0$ falls in the same bin (two-sided empirical p-value, and BH-FDR over tests). For the human lung cancer sample, the SCI-based analysis involved 4,180 protein–metabolite pairs. Because this relatively small number of pairs would otherwise yield a limited null sample size, we repeated the sampler three independent times, resulting 12,540 null SCI values to estimate empirical p-values.
